# Modulation of human dorsal root ganglion neuron excitability by Nav1.7 inhibition

**DOI:** 10.64898/2026.03.26.714428

**Authors:** Akie Fujita, Sooyeon Jo, Robert G. Stewart, Tomás Osorno, Alyssa Ferraiuolo, Kevin Carlin, Bruce P. Bean

## Abstract

Nav1.7 voltage-gated sodium channels are strongly expressed in human primary pain-sensing neurons (nociceptors) and selective Nav1.7 inhibitors have been developed as possible therapeutic agents for treating pain, so far with disappointing clinical results. In contrast, a selective Nav1.8 channel inhibitor (suzetrigine) has had successful clinical trials. Because nociceptors express both Nav1.7 and Nav1.8 channels, it is of interest to compare effects of Nav1.7 and Nav1.8 inhibitors on the excitability of human nociceptors. To compare with previous results with suzetrigine, we characterized the effects of a selective Nav1.7 inhibitor, AM-2099, on action potential generation and repetitive firing of dissociated human dorsal root ganglion neurons, studied at 37°C. Inhibition of Nav1.7 channels by 600 nM AM-2099 generally produced a substantial depolarizing shift of action potential threshold, an increase in rheobase, a decrease in action potential upstroke velocity, decrease in action potential peak, and prolongation of refractory period. Compared to inhibition of Nav1.8 channels, inhibition of Nav1.7 channels had larger effects on threshold and maximal upstroke velocity, while action potential peak was reduced similarly by both. Nav1.8 inhibition produced much more dramatic reduction of repetitive firing than Nav1.7 inhibition. The results show that although the excitability of human DRG neurons is affected by inhibition of Nav1.7 channels, most notably by an increase in threshold and increase in refractory period, repetitive firing of the neurons in response to strong stimuli is little affected. [234 words]

**Significance statement:** Nav1.7 sodium channels are highly expressed in primary pain-sensing neurons and humans with null mutations in Nav1.7 channels have loss of pain sensation. However, unlike the Nav1.8 inhibitor suzetrigine, Nav1.7 inhibitors have so far not reached clinical use. We compared effects of Nav1.7 on electrical excitability of human dorsal root ganglion neurons with those of suzetrigine and found that while Nav1.7 inhibition affects spike threshold more than suzetrigine, there is little effect on repetitive firing with strong stimuli.

## Introduction

The discovery of families with congenital insensitivity to pain traced to loss-of-function mutations in Nav1.7 voltage-dependent sodium channels (Cox et al., 2006; Goldberg et al., 2007; reviewed by Dib-Hajj et al., 2013; Drissi et al., 2020) suggested that Nav1.7 channels could be an ideal pharmacological target and launched many drug development efforts. However, PF-05089771, the first selective Nav1.7-inhibitor to reach large-scale clinical trials, was much less effective on pain in humans than anticipated (McDonnell et al., 2018; Siebenga et al., 2020; reviewed by Alsaloum et al., 2020) and many though not all Nav1.7-focused drug development efforts have been discontinued (Eagles et al., 2022; Alsaloum et al., 2025; Banh et al., 2025; Yang et al., 2025).

The reasons that Nav1.7 inhibitors have so far failed to advance to successful clinical studies are still unclear. Some compounds that are highly potent *in vitro* may have poor target engagement clinically as a result of high plasma protein binding (Deng et al., 2023). Another issue contributing to discontinuation of some Nav1.7 inhibitor development programs is observation of an acute reduction in blood pressure (Rothenberg et al., 2019; Deng et al., 2023; Mulcahy et al., 2024; Regan et al., 2024), suggesting autonomic effects reflecting expression of Nav1.7 channels in non-pain-sensing neurons (Kim et al., 2024).

In principle, one potential limitation of selectively inhibiting Nav1.7 channels is that primary pain-sensing neurons (nociceptors) express not only Nav1.7 channels but also have prominent expression of Nav1.8 channels (reviewed by Cummins et al., 2007; Bennett et al. 2019; Alles et al., 2021; Goodwin and McMahon, 2021). In the cell bodies of primary nociceptors of rat and mouse, Nav1.8 channels carry most of the sodium current driving the action potential (Renganathan et al., 2001; Blair and Bean, 2002; reviewed by Han et al., 2016) and are particularly important for supporting repetitive firing of the neurons (Elliott and Elliott, 1993; Cummins and Waxman, 1997; Rush et al., 1998; Renganathan et al., 2001; Blair and Bean, 2003; Matsutomi et al., 2006; Patrick Harty and Waxman, 2007; Han et al., 2015). Recently, a potent and highly-selective Nav1.8 inhibitor, suzetrigine, was approved by the United States Food and Drug Administration for treatment of acute pain based on results in clinical studies on post-operative pain (Jones et al., 2023; Bertoch et al., 2025). However, although suzetrigine was more effective than placebo in these studies on post-operative pain, it was far from producing complete analgesia.

We lack a good understanding of the different roles of Nav1.7 and Nav1.8 channels in controlling the excitability of human primary nociceptors. Inhibitors of both Nav1.7 and Nav1.8 channels have been shown to reduce excitability of dorsal root ganglion neurons from human donors (Payne et al., 2015; Alexandrou et al., 2016; Nguyen et al., 2022; Osteen et al., 2025; Stewart et al., 2025; Uhelski et al., 2026), but for studies of Nav1.7 inhibitors, data is limited to a handful of cells studied with limited protocols (Alexandrou et al., 2016). Studies with suzetrigine inhibition of Nav1.8 channels showed only modest effects on action potential threshold and upstroke in most human DRG neurons (Stewart et al., 2025), suggesting a possible major role of Nav1.7 channels. Here we have examined effects of a Nav1.7 inhibitor, AM-2099, on excitability of DRG neurons from human neurons. We found that Nav1.7 inhibition has larger effects than Nav1.8 inhibition on action potential threshold and action potential upstroke but much less effect on ability of the neurons to fire repetitively during stimulation by maintained depolarization. Combined with the results with suzetrigine using similar protocols, the results show that selective inhibition of either Nav1.7 or Nav1.8 channels alone has limited efficacy for inhibiting firing of human DRG neurons.

## Results

We selected AM-2099 as a suitable Nav1.7 inhibitor, based on its potency and high degree of selectivity, >200-fold for Nav1.7 over Nav1.8 (Marx et al., 2016). To evaluate the potency of the compound at 37°C, we determined its potency on human Nav1.7 channels in a stable cell line, performing concentration-dependence experiments using an automated patch clamp instrument (Sophion Qube 384) that allows temperature control (Figure 1). Interestingly, we found that AM-2099 is weaker at 37°C (IC_50_ of 109 nM) compared to 22°C (IC_50_ of 35 nM). We also tested the effect of AM-2099 on sodium channel currents in human DRG neurons on a background of suzetrigine (VX-548) to inhibit Nav1.8 channels (Figure 2). Previous voltage clamp with human DRG neurons after inhibition of Nav1.8 channels by suzetrigine showed a component of sodium current blocked by the Nav1.7 inhibiting peptide GsAF-1 together with a component of current remaining in GsAF-1 and inhibited by tetrodotoxin (TTX), with different amounts of these two components of current in different neurons (Stewart et al., 2025). Experiments applying 600 nM AM-2099 in the presence of suzetrigine showed similar results. In some neurons (e.g. Figure 2A, B), 600 nM AM-2099 inhibited almost all of the current remaining after inhibition of Nav1.8 channels by suzetrigine, with only a small additional effect of TTX. In other neurons, 600 nM AM-2099 left a larger fraction of current that was then inhibited by TTX (Figure 2C). On average, 600 nM AM-2099 inhibited 73 ± 13% of the sodium current remaining in 10 nM or 30 nM suzetrigine (Figure 2D; mean ± SD, n=10). In some neurons, we successively applied 100 nM AM-2099 then 600 nM AM-2099. In these neurons, 100 nM AM-2099 inhibited 64 ± 20% of the current inhibited by 600 nM AM-2099 (n=7). Overall, these results are consistent with an IC_50_ of roughly 100 nM for AM-2099 inhibition of native Nav1.7 channels in human DRG neurons at 37°C, consistent with the concentration-dependence data on cloned channels at 37°C (Figure 1B). Based on these results, 600 nM AM-2099 is expected to inhibit about 85% of the Nav1.7 current at 37°C. The results are also consistent with previous results with GsAF-1 in showing that TTX-sensitive sodium current in human DRG neurons is composed primarily of Nav1.7-mediated current but that some cells also have a component of non-Nav1.7 current that is inhibited by TTX (Stewart et al., 2025).

**Figure 1.**
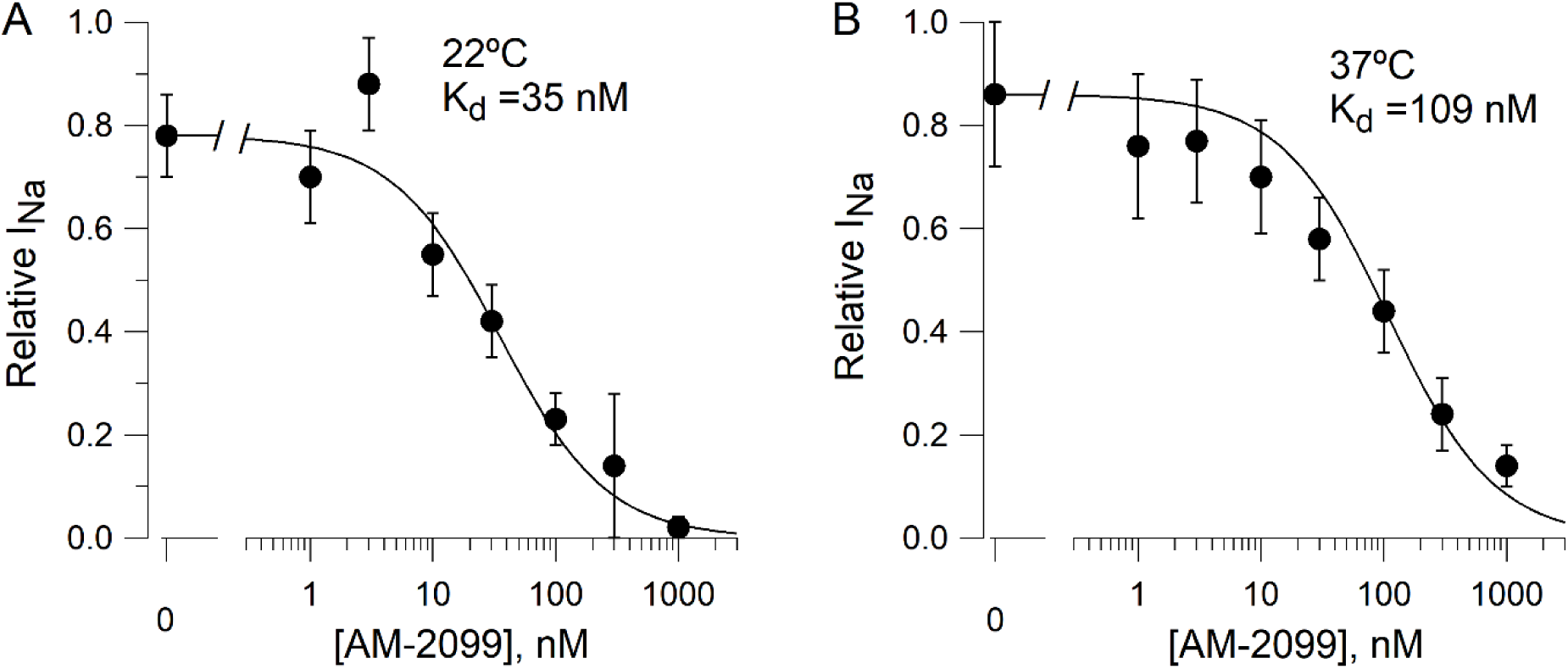
Concentration-dependence for inhibition of cloned human Nav1.7 channels by AM-2099 at 22 °C (A) and 37 °C (B). Recordings were made on a Qube 384 automated patch clamp instrument (Sophion Bioscience) using a stable cell line with human Nav1.7 channels expressed in HEK293 cells (Liu et al., 2012) using 10-hole recording wells. Sodium channel current was evoked by a step from-80 mV to +5 mV delivered every 3 seconds. Each concentration of AM-2099 was applied in 16 wells and data from each well were accepted if the leak-corrected current at the holding voltage of-80 mV was less than 0.5 nA and the peak sodium current in control was > 10 nA. Symbols show mean ± SD for wells meeting these criteria with current measured after 16 minutes of exposure to compound. A:12 wells for control solution containing 0.1% DMSO (0 nM), 8 for 1 nM, 8 for 3 nM, 8 for 10 nM, 8 for 30 nM, 8 for 100 nM, 12 for 300 nM, 15 for 1000 nM. B: 23 wells for control drug-free solution containing 0.1% DMSO, 12 for 1 nM, 13 for 3 nM, 15 for 10 nM, 9 for 30 nM, 11 for 100 nM, 11 for 300 nM, 15 for 1000 nM. Fitted curves: Imax/(1+[Drug]/Kd), where Imax was fixed at the average value in the DMSO control wells (concentration of 0 nM) and the fit included weighting of data points by SD. Fit to data in A ignored the values from the column corresponding to 3 nM AM-2099 as it was obvious that there was an error in preparing this dilution. Internal solution (mM):140 CsF, 10 NaCl, 1 EGTA, 10 HEPES, 25 sucrose pH 7.3 w CsOH. External Solution (mM):145 NaCl, 4 KCl, 2 CaCl_2_, 1 MgCl_2_, 10 HEPES, 10 Glucose, pH 7.4 w NaOH. All solutions contained 1mg/mL Pluronic-F68.

**Figure 2.**
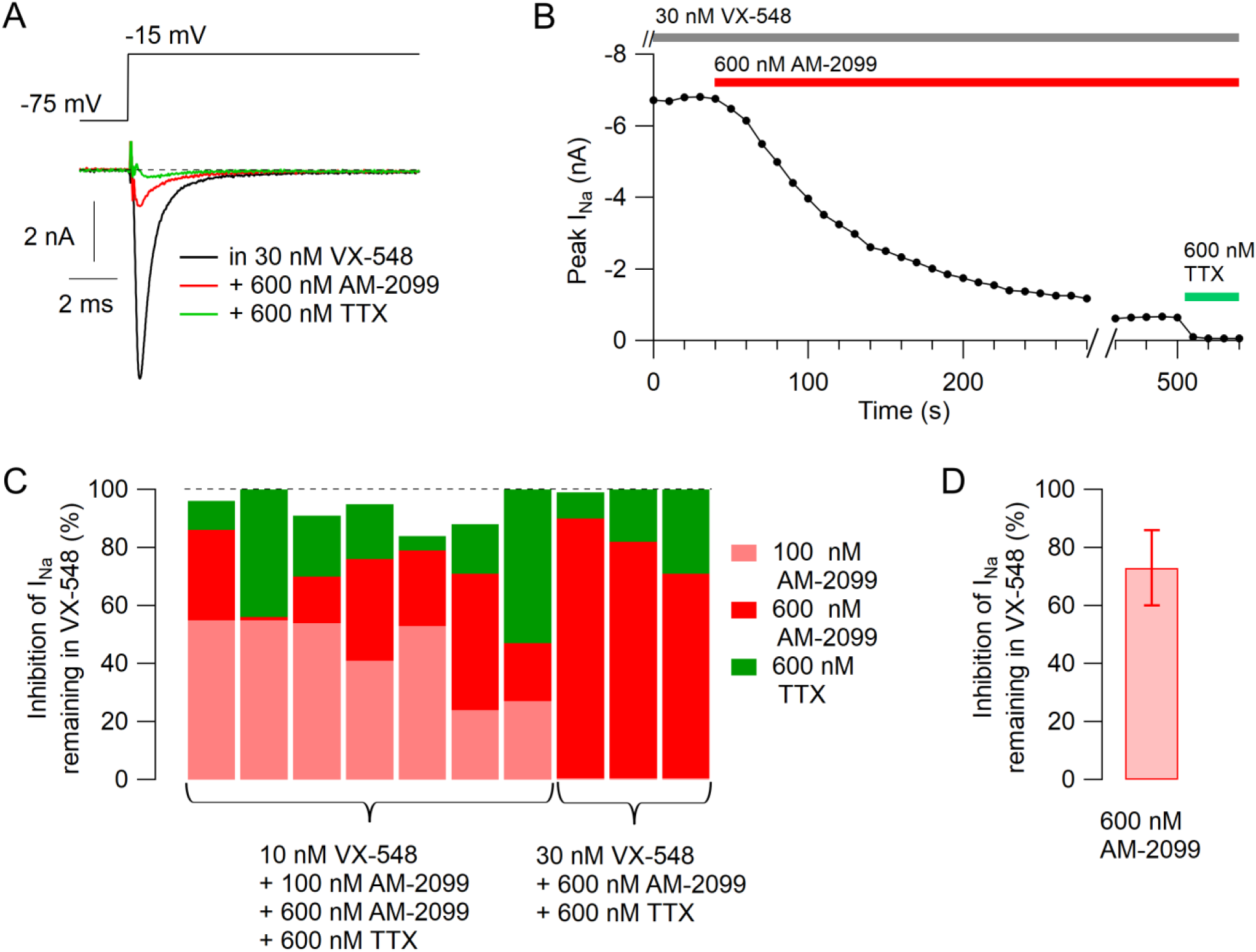
Effect of AM-2099 on voltage-clamped sodium current in human DRG neurons. A-B, Effect of cumulative addition of 600 nM AM-2099 and 600 nM TTX on sodium current evoked after the inhibition of Nav1.8 channels by 30 nM VX-548 (in the continuing presence of VX-548). C, Collected results. In 7 neurons, the cell was exposed first to 100 nM AM-2099 and then to 600 nM AM-2099 and then to 600 nM AM-2099 plus 600 nM TTX, all on a background of 10 nM VX-548. Light red bars show the fraction of current inhibited by 100 nM AM-2099, dark red bars show the additional effect of 600 nM AM-2099, and green bars show the additional effect of 600 nM TTX. In three neurons, cells were exposed to 600 nM AM-2099 and then 600 nM AM-2099 plus 600 nM TTX in the presence of 30 nM VX-548. D, Mean ± SD for the effect of 600 nM AM-2099 on the sodium current remaining in VX-548 in these ten neurons.

Figure 3 shows the effects of 600 nM AM-2099 on action potentials in human DRG neurons studied at 37°C. Action potentials were evoked by short (0.5-ms) current injections. With this protocol, the action potential shape is not affected by the stimulating current, making it easier to interpret changes in action potential shape produced by channel inhibition. Figure 3A-B shows a typical effect of AM-2099: a slowing of the rising phase of the action potential and substantially reduced peak, with little effect on the falling phase or after-hyperpolarization. Figure 3C-G show collected results for the effect of 600 nM AM-2099 on action potential parameters. In collected results, 600 nM AM-2099 produced a decrease in maximum upstroke velocity to an average of 53 ± 26% of control, from an average of 294 ± 160 to 149 ± 101 mV/ms (mean ± SD, n=40; p=3.6 x 10^-12^, two-tailed Wilcoxon test), and a decrease in action potential peak by an average of 11.2 ± 10.5 mV, from +33 ± 12 mV to +22 ± 18 mV (mean ± SD, n=40; p=0.003, two-tailed Wilcoxon test). Unlike the consistent substantial decrease in the width of the action potential seen with suzetrigine inhibition of Nav1.8 channels (Stewart et al., 2025), inhibiting Nav1.7 channels by AM-2099 had more variable effects on action potential width, decreasing the width measured at 0 mV in 24 of 34 neurons and increasing the width in 10 of 34 neurons (taking into account only neurons in which the action potential peak after AM-2099 was more positive than 0 mV). The decrease in width measured at 0 mV often likely reflected the decrease in the peak of the action potential (e.g. the example neuron in Figure 3A) rather than the clear inhibition of the action potential shoulder that was seen with suzetrigine inhibition of Nav1.8 channels (Stewart et al., 2025).

**Figure 3.**
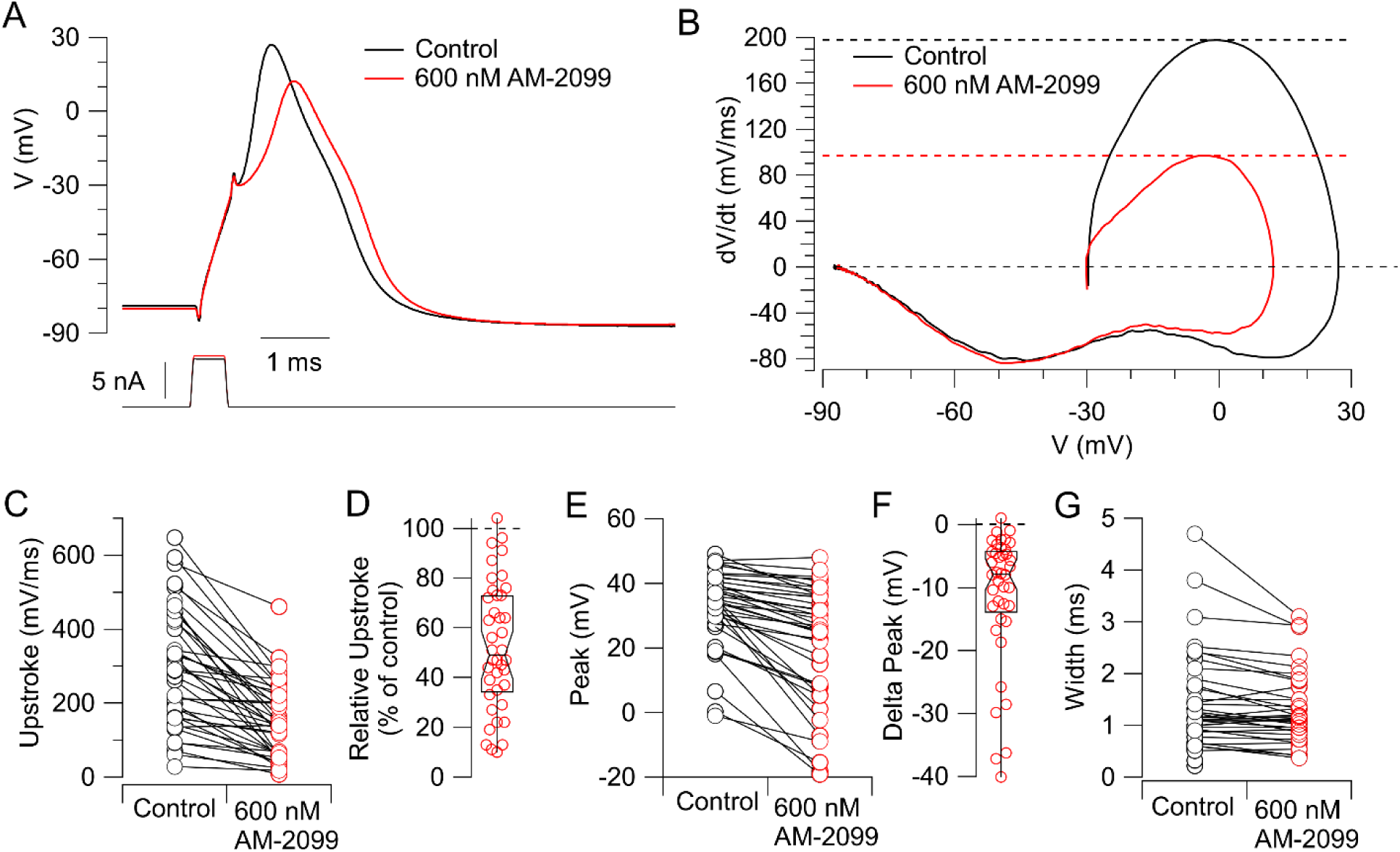
Effect of AM-2099 on action potentials in human DRG neurons. A, Action potentials in an example capsaicin-sensitive human DRG neuron evoked by a short (0.5-ms) current injection before and after application of 600 nM AM-2099. Stimulating current was adjusted to be slightly larger in AM-2099 so that the voltage immediately after cessation of the stimulating current was the same as in control (because control has an active component in the last 0.1 ms of the stimulating pulse that is inhibited with AM-2099). B, Phase-plane plot of dV/dt versus V for the action potentials in A showing reduction of maximum upstroke velocity and peak. Upper dashed lines: maximum upstroke velocity in control (black) and with 600 nM AM-2099 (red). C, Collected results for effect of 600 nM AM-2099 on the maximal upstroke velocity of the action potential in 40 neurons. D, Tukey box-plot of collected results for upstroke in 600 nM AM-2099 relative to the control. Middle bar shows median, with notch indicating 95% confidence interval for median, lower bar indicates 25% percentile, and upper bar indicates 75% percentile. E, Collected results for effect of 600 nM AM-2099 on peak of the action potential evoked by a short current injection (N=40). F, Tukey box-plot of collected results for change in action potential peak by 600 nM AM-2099. G, Collected results for effect of 600 nM AM-2099 on the width of the action potential measured at 0 mV. N=39 for cells with control peak >0 mV; values only in control shown for 5 cells in which the action potential in AM-2099 had a peak < 0 mV.

AM-2099 almost always (37 of 38 neurons) produced a depolarizing shift in the action potential threshold, determined by short (0.5-ms) current injections of increasing magnitude (Figure 4). In collected results, the threshold increased by an average of 10.5 ± 6.2 mV from-44.6 ± 8.8 mV to-34.2 ± 10.5 mV (mean ± SD, n=38; p=1.5 x 10^-11^, two-tailed Wilcoxon test). The 0.5-ms current injection required to reach threshold increased by an average of 29 ± 18% (mean ± SD, n=38; p=1.8 x 10^-7^, two-tailed Wilcoxon test).

**Figure 4.**
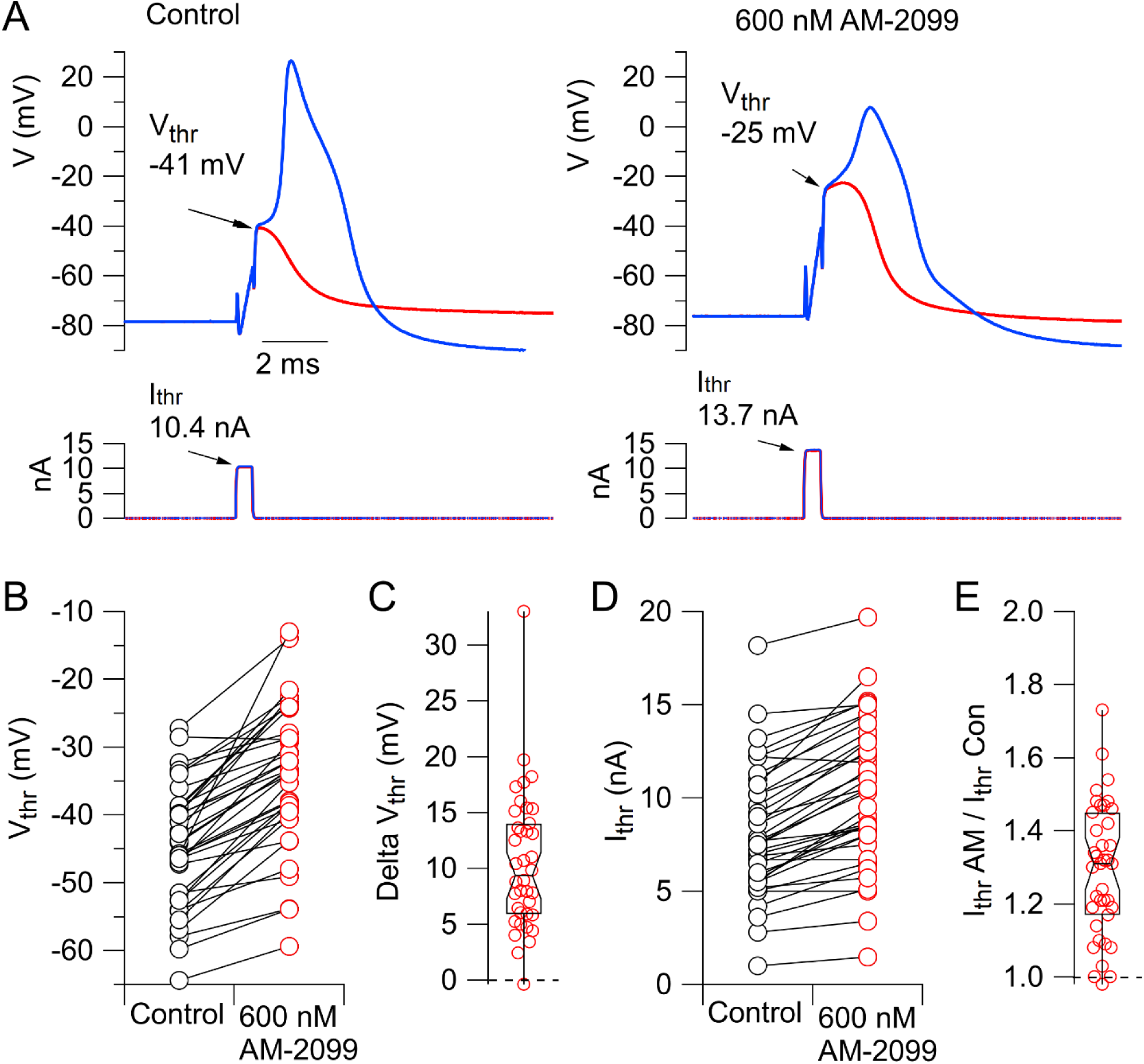
Effect of AM-2099 on action potential threshold. Short (0.5-ms) current injections of different size were delivered to find the level of current injection that first evoked an action potential. The voltage immediately after the smallest current injection that evoked a spike was considered as the threshold voltage. A, Just sub-threshold (red) and supra-threshold (blue) current injections in control (left) and after application of 600 nM AM-2099 (right). B, Collected data for threshold voltage before and after 600 AM-2099 (n=38). C, Tukey box-plot of collected results for change in voltage threshold in 600 nM AM-2099 relative to the control. Middle bar shows median, with notch indicating 95% confidence interval for median, lower bar indicates 25% percentile, and upper bar indicates 75% percentile. D, Collected data for minimum current (0.5-ms current injection) that evoked an action potential (n=38). E, Tukey box-plot of collected results of ratio of minimum current to evoke an action potential after application of 600 nM AM-2099 relative to control.

We next explored the effect of inhibiting Nav1.7 channels on repetitive firing evoked by 1-s current injections. Figure 5A-B shows a typical example: the minimal current required to elicit a spike (rheobase) increased substantially but the frequency of firing evoked by large current injections did not change much. In collected results (Figure 5C), rheobase current increased by 82 ± 68%, from 1.8 ± 1.3 nA to 3.3 ± 2.6 nA (mean ± SD, n=44; p=1.7 x 10^-7^, two-tailed Wilcoxon test). In 44 neurons tested with 1-s current injections, 31 fired multiple action potentials in control, and 29 of the 31 still fired multiple action potentials after application of AM-2099 (Figure 5D).

**Figure 5.**
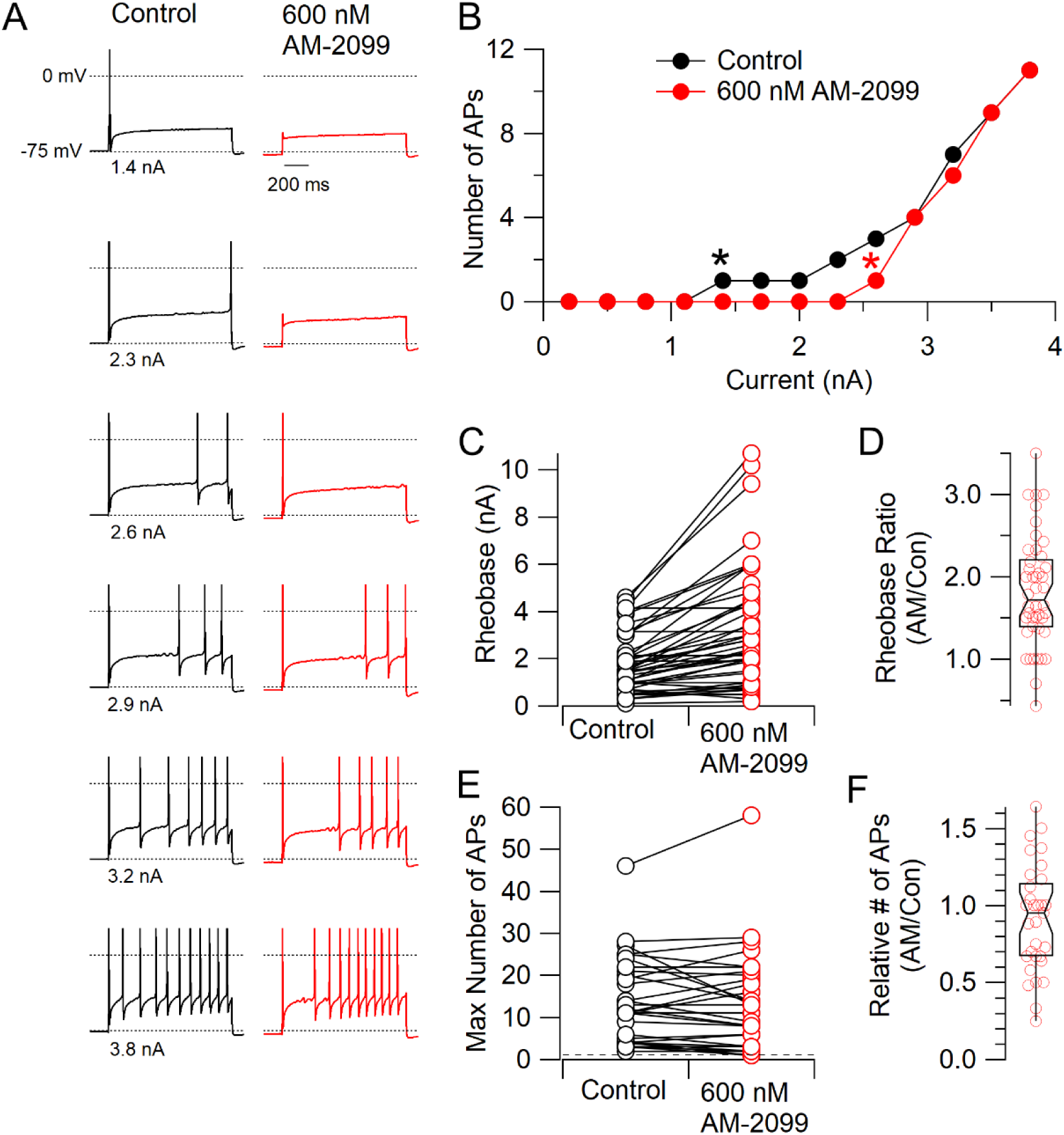
Effect of 600 nM AM-2099 on rheobase and repetitive firing. A, Firing evoked in a capsaicin-sensitive human DRG neurons by 1-s injections of current of increasing magnitude in control and after application of 600 nM AM-2099. B, Number of action potentials as a function of the injected current before and after 600 nM AM-2099 in this neuron. Asterisks indicate rheobase current in control (black) and in AM-2099 (red). C, Collected data for the effect of 600 nM AM-2099 on rheobase current. N=44 neurons. D, Tukey box-plot of collected results for rheobase in 600 nM AM-2099 relative to control rheobase. Middle bar shows median, with notch indicating 95% confidence interval for median, lower bar indicates 25% percentile, and upper bar indicates 75% percentile. E, Collected results for the effect of 600 nM AM-2099 on the maximal number of action potentials during 1-s current injections over a range of magnitudes for neurons that fired more than one action potential in control. N=31 neurons. Dashed line drawn at 1 action potential. F, Tukey box-plot of collected results for maximum number of action potentials in 600 nM AM-2099 relative to maximum number in control.

In a previous study, we were surprised to find that inhibiting Nav1.8 channels with suzetrigine had the counter-intuitive effect of decreasing the refractory period (Stewart et al., 2025). In contrast, inhibiting Nav1.7 channels resulted in an increase in the refractory period in 14 of 15 neurons tested (Figure 6), with an average increase to 1.8 ± 0.7 of the value in the control (mean ± SD, n=15; p=.0012, two-tailed Wilcoxon test).

**Figure 6.**
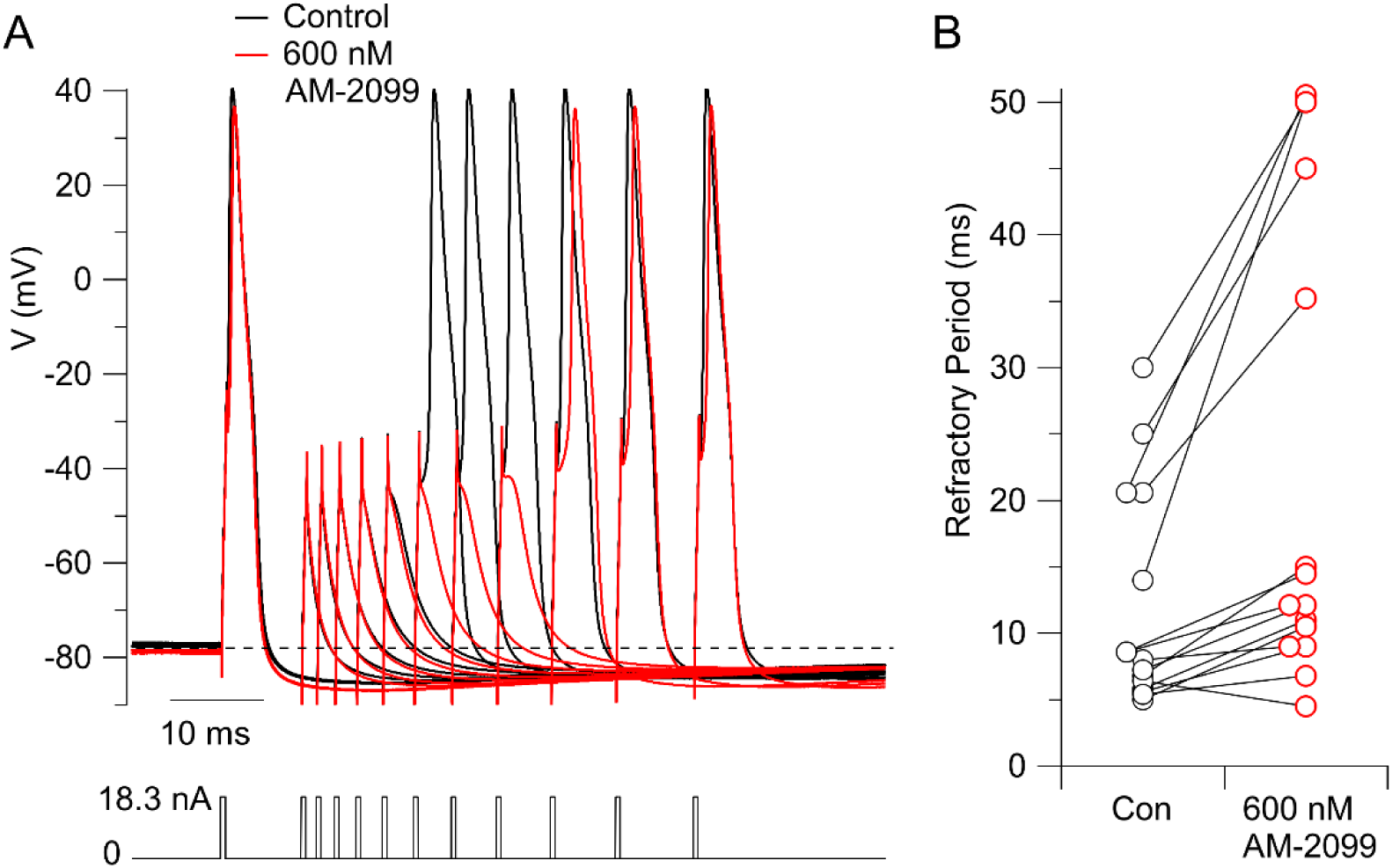
Prolongation of refractory period by AM-2099. A, Action potentials were evoked by a pair of 0.5-ms current injections with a variable time between them, with magnitude of both set at 1.5-times the threshold current determined in control. The time between the two current injections was varied from longer to shorter to determine the refractory period with this stimulus. The figure shows superimposed sweeps from 11 different sets of times in control (black) and with 600 nm AM-2099 (red). In control, the second stimulus evoked an action potential with a spacing (start to start) of 20.6 ms, while after AM-2099, a spacing of 35.2 ms was required to evoke an action potential using the same stimuli. B, Collected results in 15 neurons. In three other neurons, after application of 600 nM AM-2099 the neuron did not fire with injection of 1.5 times the threshold current in control and data are not included.

### Effects of PF-04856264

Before doing the experiments using AM-2099, we did an initial series of experiments using a different Nav1.7 inhibitor, PF-04856264 (McCormack et al., 2013). Based on the IC_50_ of 28 nM originally determined (McCormack et al., 2013), we choose a concentration of 200 nM PF-04856264, expected to produce ∼ 90% inhibition of Nav1.7 channels with minimal effects on other types of channels. In current clamp experiments, we found that 200 nM PF-04856264 decreased action potential peak, decreased maximal upstroke velocity, and shifted the threshold voltage positive (Figure S3A-E). However, quantitatively, all of these effects were somewhat smaller than the effects of 600 nM AM-2099 tested in subsequent experiments. Because of these quantitative differences, we did experiments on cloned human Nav1.7 channels to directly compare the degree of inhibition by 200 nM PF-04856264 and 600 nM AM-2099, doing experiments at 37 °C with a steady holding potential of-75 mV to match typical resting potentials of the human DRG neurons. These experiments showed that under these conditions, 200 nM PF-04856264 inhibited cloned human Nav1.7 channels by an average of 67 ± 20% (mean ± SD, n=5) and was less effective than 600 nM AM-2099, which inhibited by an average of 95 ± 4 %, (mean ± SD, n=4; Figure S3F-G). Similarly, in voltage clamp experiments in human DRG neurons, 200 nM PF-04856264 inhibited the sodium current remaining after suzetrigine inhibition of Nav1.8 channels by an average of 51 ± 25% (mean ± SD, n=9), less than the effect of 600 nM AM-2099 (average inhibition by 73 ± 13%, mean ± SD, n=10; Figure S3H). Thus, PF-04856264 had qualitatively similar effects on action potential generation as AM-2099, but at 200 nM produced only incomplete inhibition of Nav1.7 current at 37 °C. It seems likely that the reduced potency at 37 °C compared to 22 °C that we found for AM-2099 (Figure 1) may be a general property of other Nav1.7 inhibitors that inhibit by a similar mechanism of binding tightly to the domain IV voltage sensor region (Ahuja et al., 2015; McCormack et al., 2013; Kschonsak et al., 2023; Huang et al., 2024). This may be an important consideration in evaluating *in vivo* effects of such inhibitors considering that virtually all previous determinations of potency for channel inhibition have been done at room temperature.

## Discussion

Previous work has shown that, like mouse and rat small-diameter dorsal root ganglion neurons (reviewed by Rush et al., 2007; Bennett et al., 2019; Goodwin and McMahon, 2021), human dorsal root ganglion neurons express both Nav1.7 and Nav1.8 channels (Payne et al., 2015; Alexandrou et al., 2016; Zhang et al., 2017; Osteen et al., 2025; Uhelski et al., 2026). In a previous paper, we used the selective Nav1.8 inhibitor suzetrigine (VX-548) to explore how inhibition of Nav1.8 channels affects action potential generation in the neurons (Stewart et al., 2025). The results presented here using similar protocols show that inhibiting Nav1.7 channels also strongly affects the excitability of human DRG neurons but in different ways than inhibiting Nav1.8 channels. Figure 7 compares the effects of inhibiting Nav1.7 channels on action potential threshold, shape, and ability to fire repetitively with the effects of Nav1.8 inhibition determined in a previous study (Stewart et al., 2025), with recordings in both cases done at 37°C with the same protocols. Inhibiting Nav1.7 channels generally produces a larger effect on action potential threshold (Figure 7A) and maximum upstroke (Figure 7B) than Nav1.8 inhibition, while the effects on action potential peak were similar (Figure 7C). A major difference is that inhibiting Nav1.8 channels was generally much more effective than Nav1.7 inhibition in reducing repetitive firing during 1-sec current injections (Figure 7D).

**Figure 7.**
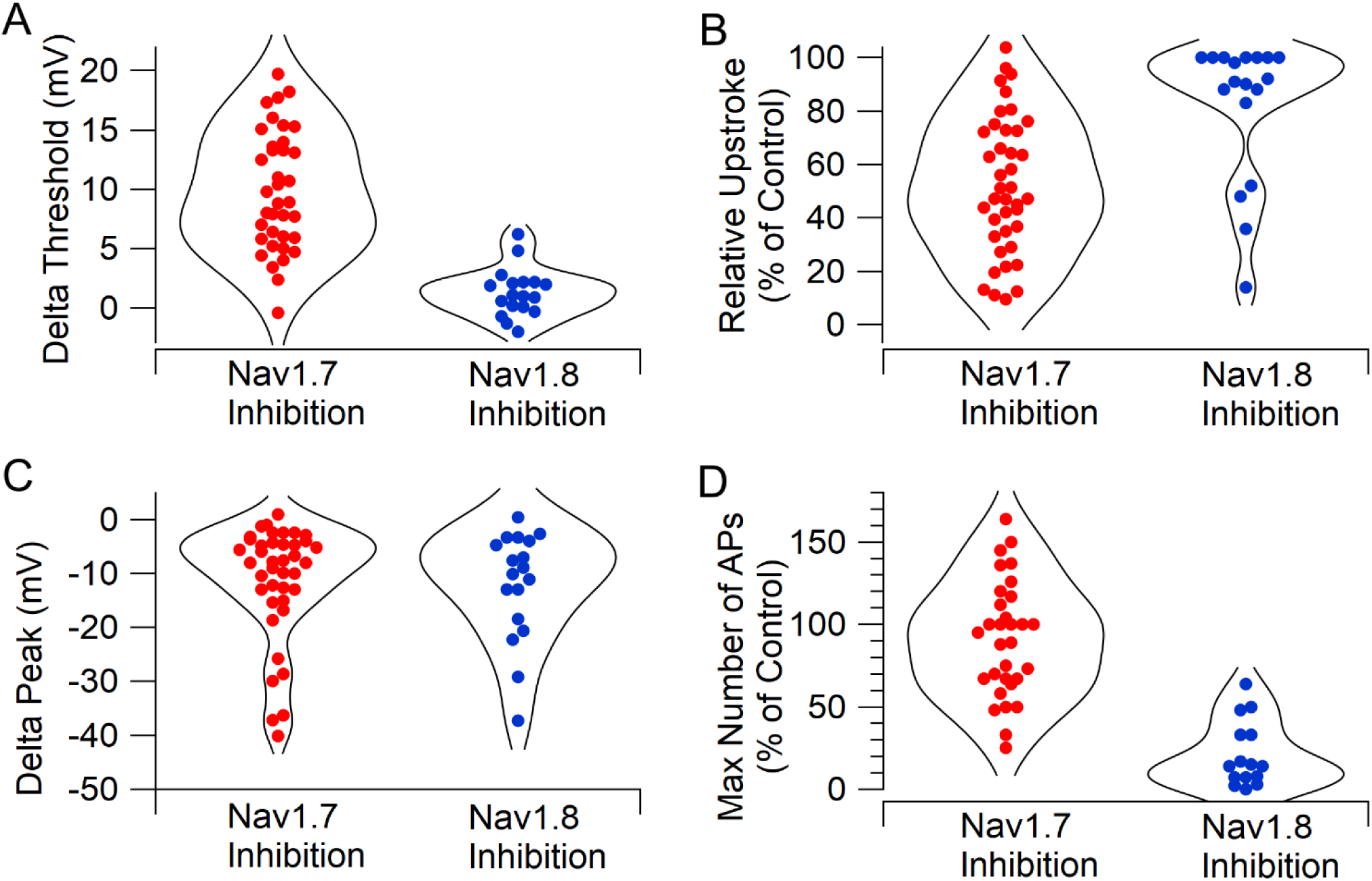
Comparison of Nav1.7 and Nav1.8 inhibition on action potential firing in human DRG neurons. A, Change in action potential threshold by inhibiting Nav1.7 channels (600 nM AM-2099, n=41 neurons) or Nav1.8 channels (10 nM VX-548, n=18 neurons). B, Inhibition of maximum upstroke of action potential by inhibiting Nav1.7 channels (600 nM AM-2099, n=41 neurons) or Nav1.8 channels (10 nM VX-548, n=18 neurons). C, Reduction of action potential peak by inhibiting Nav1.7 channels (600 nM AM-2099, n=41 neurons) or Nav1.8 channels (10 nM VX-548, n=18 neurons). D, Effect of inhibiting Nav1.7 channels (600 nM AM-2099, n=31 neurons) or Nav1.8 channels (100 nM VX-548, n=15 neurons) on maximum number of action potentials evoked by 1-s depolarizations of different magnitudes, confining analysis to cells that fired more than one action potential in control. Data for effects of VX-548 from Stewart et al., (2025).

The different effects of inhibiting Nav1.7 and Nav1.8 channels on action potential shape and repetitive firing fit well with their different voltage-dependence and kinetics as defined from studies on both human and rodent channels (Alexandrou et al., 2016; Köster et al., 2025; Vasylyev et al., 2024, 2025) and with the different but overlapping roles of Nav1.7 and Nav1.8 channels in action potentials of mouse and rat DRG neurons (Renganathan et al., 2001; Blair and Bean, 2002; Matsutomi et al., 2006; Rush et al., 2007; Han et al., 2015a; Shields et al., 2018; Vasylyev et al., 2024, 2025; Xie et al., 2024). Nav1.7 channels are activated more quickly and at more negative voltages than Nav1.8 channels, consistent with the dominant effect of Nav1.7 channels on threshold and maximum upstroke velocity. Nav1.8 channels inactivate more slowly than Nav1.7 channels and a substantial fraction of Nav1.8 channels remain open during the shoulder of the action potential (Stewart et al., 2025). An apparent quantitative difference between human and rodent DRG neurons is the dominance of Nav1.7 channels in the upstroke of the action potential in human neurons, while TTX-resistant current from Nav1.8 channels appears to dominate in the upstroke of action potentials in small-diameter rat and mouse neurons (Renganathan et al., 2001; Blair and Bean, 2002; Matsutomi et al., 2006; Rush et al., 2007; Han et al., 2015a). Interestingly, this quantitative difference between mouse and rat DRG neurons was mirrored in a PCR analysis of seven sodium channel subtypes in DRG neurons of human and mouse, showing higher relative expression of Nav1.7 compared to Nav1.8 in human neurons compared to mouse neurons (Chang et al., 2018).

Our results are generally consistent with those of Alexandrou et al. (2016), who found that the Nav1.7 inhibitor PF-05089771 blocked action potential generation in most but not all (8 of 15) human DRG neurons tested when action potentials were evoked by current injections that were just-suprathreshold in control. We saw more consistent effects of inhibiting Nav1.7 channels, with an increase in voltage threshold in 14 of 15 neurons. The difference could reflect in part the different recording conditions (37°C in our experiments versus room temperature in the Alexandrou et al. recordings) but could also reflect intrinsic heterogeneity in cell populations.

The strong effect of inhibiting Nav1.7 current on action potential threshold is reflected in the increase in rheobase current necessary to evoke spiking in the 1-s current injection protocols (Figure 5). The effect of inhibiting Nav1.7 channels with AM-2099 is concordant with an increase in rheobase in iPSC-derived nociceptors from human patients with Nav1.7 loss-of-function mutations (McDermott et al., 2019) and with an increase in rheobase with the Nav1.7 inhibitor PF-05089771 in iPSC neurons from healthy controls but not patients with Nav1.7 loss-of-function mutations (McDermott et al., 2019).

In contrast to the counter-intuitive decrease in refractory period seen with Nav1.8 inhibition (Stewart et al., 2025), Nav1.7 inhibition almost always (14 of 15 neurons) increased the refractory period. The overall effects on refractory period of inhibiting either channel reflect a complex combination of the actual reduction in sodium channel availability along with secondary effects resulting from the change in action potential shape. The increase in refractory period with Nav1.7 inhibition is easily understood from the fact that Nav1.7 channels strongly control action potential threshold. The reduction in action potential peak (seen with both Nav1.7 and Nav1.8 inhibition) and a reduction in action potential width (which was dramatic with Nav1.8 inhibition but generally not with Nav1.7 inhibition) would both be expected to have a secondary effect of tending to reduce the refractory period by producing less activation of voltage-activated potassium channels and therefore reducing the post-spike potassium conductance that contributes to the refractory period. Because inhibiting Nav1.8 channels has much less effect on action potential threshold compared to inhibiting Nav1.7 channels, the effects on potassium conductance to decrease refractory period – resulting from the changes in action potential shape - apparently dominate with Nav1.8 inhibition but not Nav1.7 inhibition.

A striking aspect of the data is the high degree of neuron-to-neuron variability in the effects of inhibiting Nav1.7 channels. Although the median effect of AM-2099 was to reduce maximum upstroke velocity by 47%, there were rare cells where upstroke velocity was reduced very little (4 of 40 cells with reduction by <10%) and other cells where upstroke velocity was reduced very dramatically (4 of 40 cells with reduction by >85%) (Figure 3D). Clearly, there are large neuron-to-neuron differences in the relative expression of Nav1.7 versus other sodium channels, even in in this population of DRG neurons verified to be capsaicin-sensitive. In the future, larger data sets in combination with single cell measurements of mRNA levels may help interpret the large cell-to-cell variability in terms of defined subpopulations of C-fiber nociceptors (cf. Shiers et al., 2020; Middleton et al., 2021; Tavares-Ferreira et al., 2022; Körner et al., 2022, 2026). Small-diameter mouse DRG neurons also have a large degree of neuron-to-neuron variability in the relative magnitude of Nav1.7 and Nav1.8 currents, even when experiments are restricted to CGRP-lineage capsaicin-sensitive mouse DRG neurons (Osorno et al., 2026). An interesting possibility is that neurons with different combinations of Nav1.7 and Nav1.8 channels could be involved in different pain states (Vasylyev et al., 2026), which could help explain why some types of pain are more sensitive to Nav1.7 or Nav1.8 knock-out or inhibition than others (Shields et al., 2018; reviewed by Yang et al., 2025).

A limitation of our experiments is that the 600 nM concentration of AM-2099 that we used produced an estimated ∼85% rather than complete inhibition of the Nav1.7 current in the neurons. We choose the concentration of 600 nM to minimize any chance that effects of the compound might include inhibition of non-Nav1.7 components of current in the neurons. Although the selectivity for AM-2099 is >200-fold for Nav1.7 over Nav1.8 channels, the selectivity is less (15-26-fold) over Nav1.2 and Nav1.6 channels (Marx et al., 2016), which could contribute to the component of current in the human DRG neurons that is inhibited by tetrodotoxin but resistant to Nav1.7 inhibitors (Stewart et al., 2025; Figure 2 of this paper). Further characterization of concentration-dependence and selectivity of AM-2099 or other Nav1.7 inhibitors would be useful to identify concentrations optimal for maximal selective inhibition of Nav1.7 channels. An important consideration for such characterization - and for interpreting *in vivo* effects of inhibitors - is the weaker potency at 37°C compared to room temperature that we saw for the action of AM-2099 on Nav1.7 channels (Figure 1).

Experiments on neuronal cell bodies like those reported here can help define the effects of sodium channel inhibitors on channels controlling human neuron excitability but may be of limited value for predicting clinical effects on pain. A major limitation in our current knowledge of nociceptor excitability is very limited information about the electrophysiology of the key functional elements of the neurons: the peripheral terminals where action potentials are first generated, the axons where the action potentials are propagated, and the fine branches and synaptic boutons of the central terminals where action potentials trigger synaptic transmission (Bennett et al., 2019; Goodwin and McMahon, 2021). Nav1.7 channels are expressed in all of these regions (Black et al., 2012; Tyagi et al., 2025) but their electrophysiological roles relative to Nav1.8 or other sodium channels in the various compartments remain to be well-defined and may vary among neuronal types. Nav1.7 channels are clearly involved in generation and propagation of action potentials in some neurons, because selective Nav1.7 inhibitors applied to the nerve can inhibit firing of C-fibers evoked by cutaneous stimulation, although often not completely (Goodwin et al., 2022; Deng et al., 2023). A study using microneurography showed that high doses of a Nav1.7 inhibitor delivered intravenously to rhesus monkeys inhibited propagating action potentials evoked by cutaneous stimulation completely in 7 of 8 nociceptive C-fibers (Kraus et al., 2021), contrasting with much weaker effects seen subsequently with a Nav1.8 inhibitor (Vardigan et al., 2025). In mice and pigtail monkeys, C-fiber action potentials are inhibited to a greater extent by tetrodotoxin in dorsal roots and proximal nerves than in distal peripheral branches, where Nav1.8 channels can support tetrodotoxin-resistant conduction to at least some extent (Klein et al., 2017), suggesting differential functional expression of Nav1.7 and Nav1.8 channels in different locations. A recent study making *in vivo* pig single nerve fiber recordings of C-fiber nociceptors excited by transcutaneous stimulation found that inhibiting Nav1.7 channels by protoxin II reduced firing more dramatically at low stimulus intensities than high intensities (Soares et al., 2025), which could be consistent with Nav1.7 channels regulating firing near threshold stimulation but Nav1.8 channels being more important for repetitive firing with strong stimulation.

While recognizing the limitations of studies on cell bodies, our results raise the possibility that a factor distinguishing effects of inhibiting Nav1.7 and Nav1.8 channels could be the greater effect of Nav1.8 inhibition to reduce repetitive firing of the neurons compared to Nav1.7 inhibition. It seems plausible that this difference may extend to initial generation of neuronal firing in the peripheral terminals, where computer modeling suggests that Nav1.8 channels play a major role in regulating repetitive firing (Barkai et al., 2020). The perception of pain is correlated with higher firing frequencies of nociceptors (Yarnitsky et al., 1992; Djouhri et al., 2006; reviewed by Namer and Lampert, 2025), so it is possible that the greater efficacy of Nav1.8 inhibition compared to Nav1.7 inhibition to reduce repetitive firing of human nociceptive neurons contributes to differences in clinical efficacy.

The expression of Nav1.7 channels can be up-regulated by pro-inflammatory mediators (Huang et al., 2014; Tyagi et al., 2024) and in various pathological conditions associated with pain (reviewed by Alsaloum et al., 2025; Yogi et al., 2025) including chemotherapy-induced peripheral neuropathy (Li et al., 2018; Akin et al., 2021; Braden et al., 2022) and early-life stress (Alvarez et al., 2021). The differing efficacy of various Nav1.7 inhibitors in a variety of animal models of pain (Alsaloum et al., 2025; Yogi et al., 2025) suggests that there may be particular forms of human pain that would respond to Nav1.7 inhibition more robustly than the acute pain initially tested. Further studies of roles of Nav1.7 channels in excitability of human nociceptors in conditions such as chemotherapy-induced peripheral neuropathy and other forms of human neuropathic pain (Raja et al., 2020; Stucky and Mikesell, 2021) would be illuminating and may be feasible as it becomes possible to test neurons from human donors with known pain conditions.

## Materials and Methods

### Automated patch clamp determination of AM-2099 concentration-dependence

The concentration-dependence for inhibition of cloned human Nav1.7 channels by AM-2099 was determined using a Qube 384 automated patch clamp instrument (Sophion Bioscience) using a stable cell line with human Nav1.7 channels expressed in HEK293 cells (Liu et al., 2012). Cells were grown and passaged at 37°C in 5% CO_2_ in Eagle’s Minimum Essential Medium (EMEM) (ATCC, CAT# 30-2003) supplemented with 10% fetal bovine serum (Gibco), 1% penicillin/streptomycin solution (Gibco, CAT# 15140-122) and 800 µg/mL Genetecin (G418 sulfate) (Gibco, Cat #10131-027). To prepare cells for Qube experiments, cells were seeded at 3-6 x 10^6^ in 150 mm Falcon cell culture dishes (Corning, CAT# 353025) and were passaged every 2-3 days at a confluence of 80-90%. On the day of the Qube experiments, cells were detached with 3 mL Detachin solution (Amsbio, CAT# AMS.T100110) applied at 37°C for 3 minutes. Cells were diluted with FreeStyle^TM^ CHO expression medium (ThermoFisher Gibco Cat# 12651014) in 15 mL Falcon tubes and centrifuged for 2 minutes at 65G. The cell pellet was diluted in FreeStyle^TM^ CHO expression medium, adjusted to a final cell density of 1.5-2.5 x 10^6^/mL). Cell density and viability (97%–98%) were measured with a Countess 3 cell counter (Invitrogen).

Recordings were made using 384-well QChip 384X plates with 10-hole recording wells. The internal solution was (in mM) 140 CsF, 10 NaCl, 1 EGTA, 10 HEPES, 25 sucrose pH 7.3 w CsOH and the external solution was (in mM):145 NaCl, 4 KCl, 2 CaCl_2_, 1 MgCl_2_, 10 HEPES, 10 Glucose, pH 7.4 w NaOH. All external solutions contained 1 mg/mL Pluronic-F68 (Sigma), a surfactant poloxamer that minimizes apparent loss of low concentrations of compounds by the high surface-to-volume microfluidics in automated patch clamp instruments (Vaelli et al., 2024). Sodium channel current was evoked by a step from-80 mV to +5 mV delivered every 3 seconds. After 5 minutes of recording in control solution to allow stabilization of currents, compound was applied by three successive liquid additions (18 μL each time) at intervals of 3 minutes and effects were measured 10 minutes after the final liquid addition (16 minutes of total exposure). Each concentration of AM-2099 was applied in 16 wells and data from each well were accepted if the leak-corrected current at the holding voltage of-80 mV was less than 0.5 nA and the peak sodium current in control was > 10 nA. Experiments were done with the temperature controlled to either 22°C or 37°C.

### Preparation of human DRG neurons

Neurons were obtained from the dorsal root ganglia (DRGs) of human donors. The procurement network of AnaBios Corporation includes only US-based Organ Procurement Organizations and Hospitals. Policies for donor screening and consent are those established by the United Network for Organ Sharing (UNOS). Organizations supplying human tissues to AnaBios follow the standards and procedures established by the US Centers for Disease Control (CDC) and are inspected biannually by the Department of Health and Human Services (DHHS). The distribution of donor medical information is in compliance with HIPAA regulations to protect donor privacy. All transfers of donor tissue to AnaBios are fully traceable and periodically reviewed by US Federal authorities.

Neurons were dissociated from dorsal root ganglia and suspended in medium (Davidson et al., 2014) and shipped in suspension in a container with temperature maintained at 4°C (Lesnak et al., 2025). The neurons were then plated on round 12 mm poly-D-lysine-treated coverslips placed in 24-or 48-well plates. Coverslips (Fisherbrand, Cat#12-545-80) were prepared by exposure UV for approximately 15-30 minutes and then incubated overnight at 4°C with 0.01 mg/mL poly-D-lysine diluted in sterile water. After trituration of the tissue, 60 μL of the sample was plated on each coverslip and incubated at 37°C (5% CO_2_) for 1-2 hours to allow the cells to settle and attach to the coverslip. Each well was then gently flooded with 1 mL of culture media consisting of BrainPhys media (Stemcell technologies, Cat. 05709), 1% penicillin/streptomycin, 1% GlutaMAX, 2% NeuroCult SM1 (Stemcell technologies, Cat 05711), 1% N-2 Supplement (Thermo Scientific, Cat # 17502048) and 2% Fetal Bovine Serum. Plates were housed in a 5% CO_2_ incubator set at 37°C for up to 6 days.

Table S1 gives information on age, sex, ethnicity, body mass index, and cause of death for the 7 donors whose neurons were used for the experiments, along with break-downs of how many neurons from each donor were used for data in each figure.

### Electrophysiology with human DRG neurons

For recording, coverslips were placed into the recording chamber containing about 2 mL of Tyrode’s solution at room temperature. We generally selected smaller cells for recording and selected cells in which glia had partially peeled off, which facilitated recording. Prior to recording, if the cell was strongly adhered to the coverslip (as was usual 2-3 days after plating), an electrode with a blunt tip was used to scrape the surrounding area of the cell and gently maneuvered to ensure detachment from the coverslip. Both the “scraping” electrode and recording electrode were pulled from borosilicate capillaries (VWR International, Cat #53432-921) on a Sutter P-97 puller (Sutter Instruments).

Supplementary Figure S1 shows histograms and box-plots of cell capacitance, calculated cell diameter, input resistance, resting potential, and maximum upstroke velocity of the action potential for the cells whose data was used in this study (all capsaicin-sensitive)..

Whole-cell patch-clamp recordings were made with either an Axon Instruments Multiclamp 700B amplifier (Molecular Devices) controlled by pClamp9.2 software (Axon Instruments), filtered at 10 kHz with a low-pass Bessel filter, and digitized at 100 kHz by a Digidata 1322A data acquisition interface or a Sutter Instruments dPatch integrated amplifier/data acquisition system, with filtering at 10 kHz and digitization at 100 kHz. Whole-cell recordings were obtained using patch pipettes with resistances of 1.2-2.5 MΩ. The shank of the electrode was wrapped with Parafilm (American National Can Company) in order to reduce capacitance and allow optimal series resistance compensation (bridge balance) of 60-80% without oscillation. Seals were obtained and the whole-cell configuration established with cells in room-temperature Tyrode’s solution consisting of 155 mM NaCl, 3.5 mM KCl, 1.5 mM CaCl_2_, 1 mM MgCl_2_, 10 mM HEPES, 10 mM glucose, pH adjusted to 7.4 with NaOH. After establishing whole-cell recording, cells were lifted off the bottom of the recording chamber and placed in front of an array of quartz flow pipes (250 μm internal diameter, 350 μm external diameter, Polymicro Technologies) attached with styrene butadiene glue (Amazing Goop, Eclectic Products) to an aluminum rod (1×1 cm) whose temperature was controlled by resistive heating elements and a feedback-controlled temperature controller (TC-344B; Warner Instruments). Solution changes were made (in < 1 second) by moving the cell between adjacent pipes. Experiments were done with temperature controlled at 37°C.

### Current clamp recordings from human DRG neurons

Current clamp recordings were made using patch pipettes filled with a K-gluconate-based internal solution containing (in mM) 139.5 K-Gluconate, 1.6 MgCl_2_, 1 EGTA, 0.09 CaCl_2_, 9 HEPES,14 creatine phosphate (Tris salt), 4 MgATP, 0.3 GTP (Tris salt), pH adjusted to 7.2 with KOH. Membrane potentials are corrected for a liquid junction potential of-13 mV between the internal solution and the Tyrode’s solution in which the current was zeroed before recording.

Cells were recorded at their natural resting membrane potential without any injection of steady holding current. For the data in Figures 1, 2, and 4, action potentials were evoked by short (0.5-ms) injections of current so that the action potential occurred after the current injection. For the data in Figure 5, 1-s current steps were applied with the increments of the steps adjusted depending on the input resistance of each cell. Action potentials were defined using a criteria of action potential peak >-20 mV and height >40 mV. Experiments were included in combined data sets only if the resting potential varied by less than 5 mV during the application of AM-2099. Capsaicin sensitivity was tested at the end of the experiment by holding the cell at-70 mV in voltage clamp and briefly applying 1 μM capsaicin and data were used only from capsaicin-sensitive neurons, which comprised ∼95% of the neurons that survived to the point of being tested with capsaicin.

A notable feature of the effects of AM-2099 inhibition of Nav1.7 channels was a large degree of neuron-to-neuron variability in the quantitative effects. As shown in Supplementary Figure S2, this variability was clear even in the neurons from individual donors.

### Voltage clamp recordings from human DRG neurons

Voltage clamp recordings examining the pharmacology of sodium current in the human DRG neurons were done using an internal solution consisting of 61 mM CsF, 61 mM CsCl, 9 mM NaCl, 1.8 mM MgCl_2_, 1.8 mM EGTA, 14 mM creatine phosphate (tris salt), 4 mM MgATP, 0.3 mM GTP (tris salt), and 9 mM HEPES, pH adjusted to 7.2 with CsOH. Membrane potentials are corrected for a liquid junction potential of-5 mV between the internal solution and the Tyrode’s solution in which the current was zeroed before recording. Seven cells tested the effect of 100 nM AM-2099 followed by 600 nM AM-2099 using an external solution consisting of 77.5 mM NaCl, 77.5 mM tetraethylammonium chloride, 5 mM BaCl2, 0.03 mM CdCl2, 10 mM HEPES, 10 mM glucose, 1 mg/mL Pluronic F-68, and 10 nM VX-548. Three cells tested the effect of 600 nM VX-548 alone using an external solution consisting of 155 mM NaCl, 10 mM tetraethylammonium chloride, 5 mM BaCl2, 0.03 mM CdCl2, 10 mM HEPES, 10 mM glucose, 1 mg/mL Pluronic F-68, and 30 nM VX-548. Analyzed data were confined to neurons in which the peak current before addition of AM-2099 (in the presence of VX-548) was < 12 nA. As for current clamp experiments, data were confined to capsaicin-sensitive neurons.

### Manual patch clamp with cloned human Nav1.7 channels

Experiments comparing the effects of AM-2099 and PF-04856264 on cloned human Nav1.7 channels studied in isolation were done using the same stable cell line with human Nav1.7 channels expressed in HEK293 cells as used in the automated patch clamp experiments (Liu et al., 2012). To best match the methods of application in the patch clamp recordings from DRG neurons, these experiments were done using manual patch clamp, with solutions applied using quartz flow pipes with continual flow. The external solution was Tyrode’s solution consisting of 155 mM NaCl, 3.5 mM KCl, 1.5 mM CaCl_2_, 1 mM MgCl_2_, 10 mM HEPES, 10 mM glucose, pH adjusted to 7.4 with NaOH. Two different internal solutions were used. One was a CsCl/CsF-based internal solution consisting of 61 mM CsF, 61 mM CsCl, 9 mM NaCl, 9 mM tetraethylammonium Cl, 1.8 mM MgCl_2_, 9 mM HEPES, 1.8 mM EGTA,14 mM creatine phosphate (tris salt), 4 mM MgATP, and 0.3 mM GTP (tris salt), pH adjusted to 7.2 with CsOH. The other was the same potassium gluconate-based internal solution used for the current clamp recordings from the DRG neurons. Sodium current was activated by 5-ms depolarizations to-3 mV or +5 mV from a holding voltage of-75 mV, chosen to be similar to typical resting potentials in the experiments on DRG neurons. Test pulses were delivered every 10 seconds. There was no clear difference in the effects of the compounds using the two different internal solutions, and results were combined. Experiments were done at 37°C.

### Drugs

AM-2099 (Marx et al., 2016) and PF-04856264 (McCormack et al., 2013) were purchased from MedChemExpress (AM-2099: Catalog HY-100727, 98.4% purity; PF-04856264: Catalog HY-12811, 99.3% purity). Stock solutions were prepared at 10 mM in DMSO, aliquoted, and frozen at-20°C. The synthesis of the active enantiomer of VX-548 (98.9 % purity) is described in Vaelli et al., (2024); stocks were prepared at 10 mM in DMSO and aliquoted and frozen at-20°C. Capsaicin stock was prepared at 1 mM from powder in DMSO and stored at room temperature. All external control solutions contained concentrations of DMSO to match those in the drug-containing solutions. All external solutions contained 1 mg/mL Pluronic PF-68 (Sigma).

## Data analysis and statistics

Data were analyzed using programs written in Igor Pro 6, 8 or 9 (Wavemetrics, Lake Oswego, OR), using DataAccess (Bruxton Software) to read pClamp files into Igor Pro.

## Acknowledgements

This work was supported by National Institutes of Health Grant R35-NS127216. Funds supporting acquisition and preparation of the neurons were from the AnaBios Corporation. We are very grateful to the anonymous donors and their families for providing the human DRG neurons used in this study. We thank Richard Kondo for facilitating transfer of the preparation of the human DRG tissue.

## Author Contributions

AF, SJ, RGS, and TO designed and executed experiments, analyzed data, and contributed to writing the manuscript; AF and KC designed procedures for and supervised dissection and preparation of tissue; BPB helped design experiments, analyze data, and write the manuscript.

## Competing Interest Statement

None of the authors have competing interests.

## Supporting Information

**Table S1.**
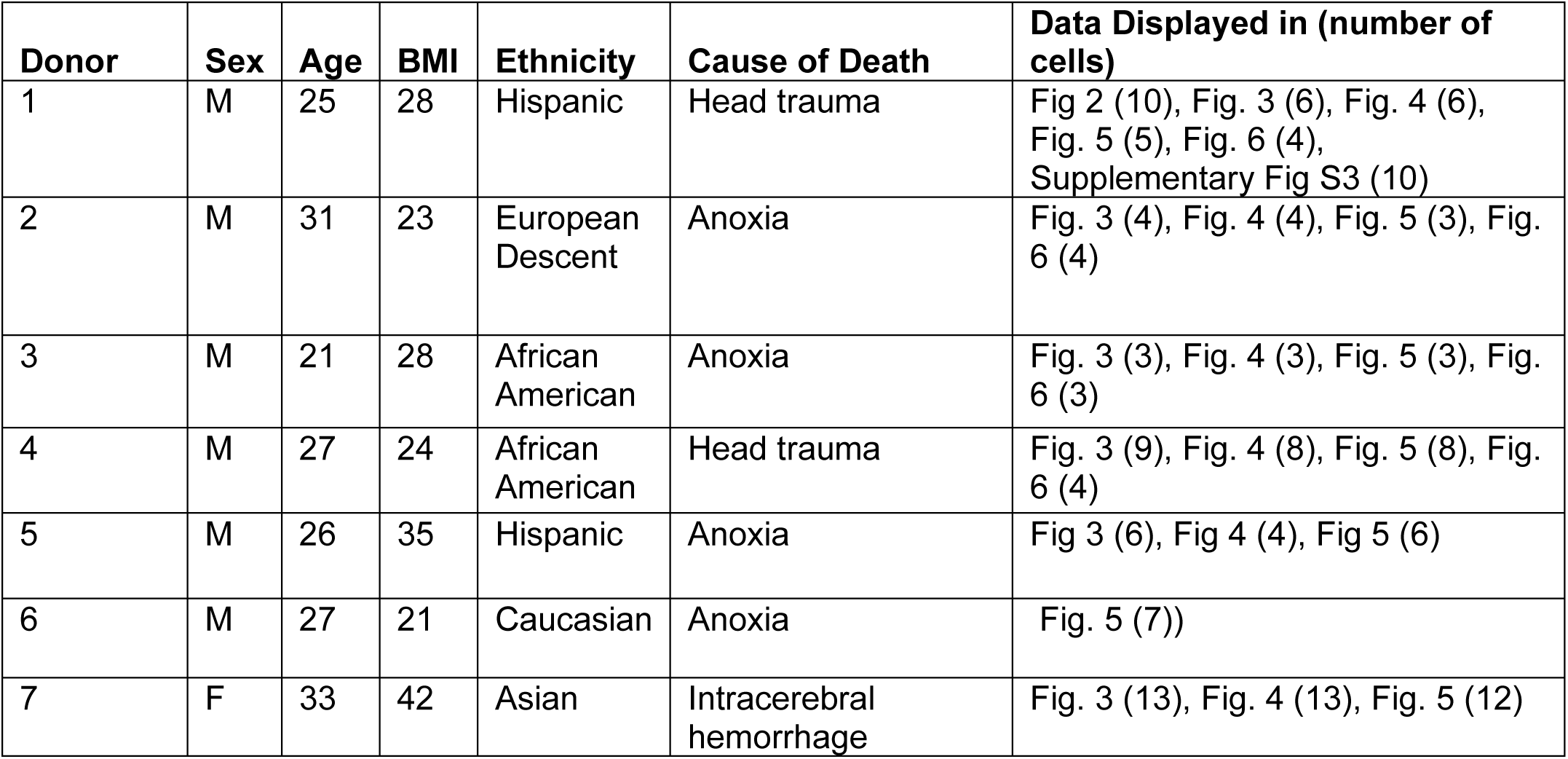
Donor demographics and distribution of data from cells used from each donor.

| **Donor** | **Sex** | **Age** | **BMI** | **Ethnicity** | **Cause of Death** | **Data Displayed in (number of cells)** |
| --- | --- | --- | --- | --- | --- | --- |
| 1 | M | 25 | 28 | Hispanic | Head trauma | Fig 2 (10), Fig. 3 (6), Fig. 4 (6), Fig. 5 (5), Fig. 6 (4), Supplementary Fig S3 (10) |
| 2 | M | 31 | 23 | European Descent | Anoxia | Fig. 3 (4), Fig. 4 (4), Fig. 5 (3), Fig. 6 (4) |
| 3 | M | 21 | 28 | African American | Anoxia | Fig. 3 (3), Fig. 4 (3), Fig. 5 (3), Fig. 6 (3) |
| 4 | M | 27 | 24 | African American | Head trauma | Fig. 3 (9), Fig. 4 (8), Fig. 5 (8), Fig. 6 (4) |
| 5 | M | 26 | 35 | Hispanic | Anoxia | Fig 3 (6), Fig 4 (4), Fig 5 (6) |
| 6 | M | 27 | 21 | Caucasian | Anoxia | Fig. 5 (7)) |
| 7 | F | 33 | 42 | Asian | Intracerebral hemorrhage | Fig. 3 (13), Fig. 4 (13), Fig. 5 (12) |

---

**Figure S1.**
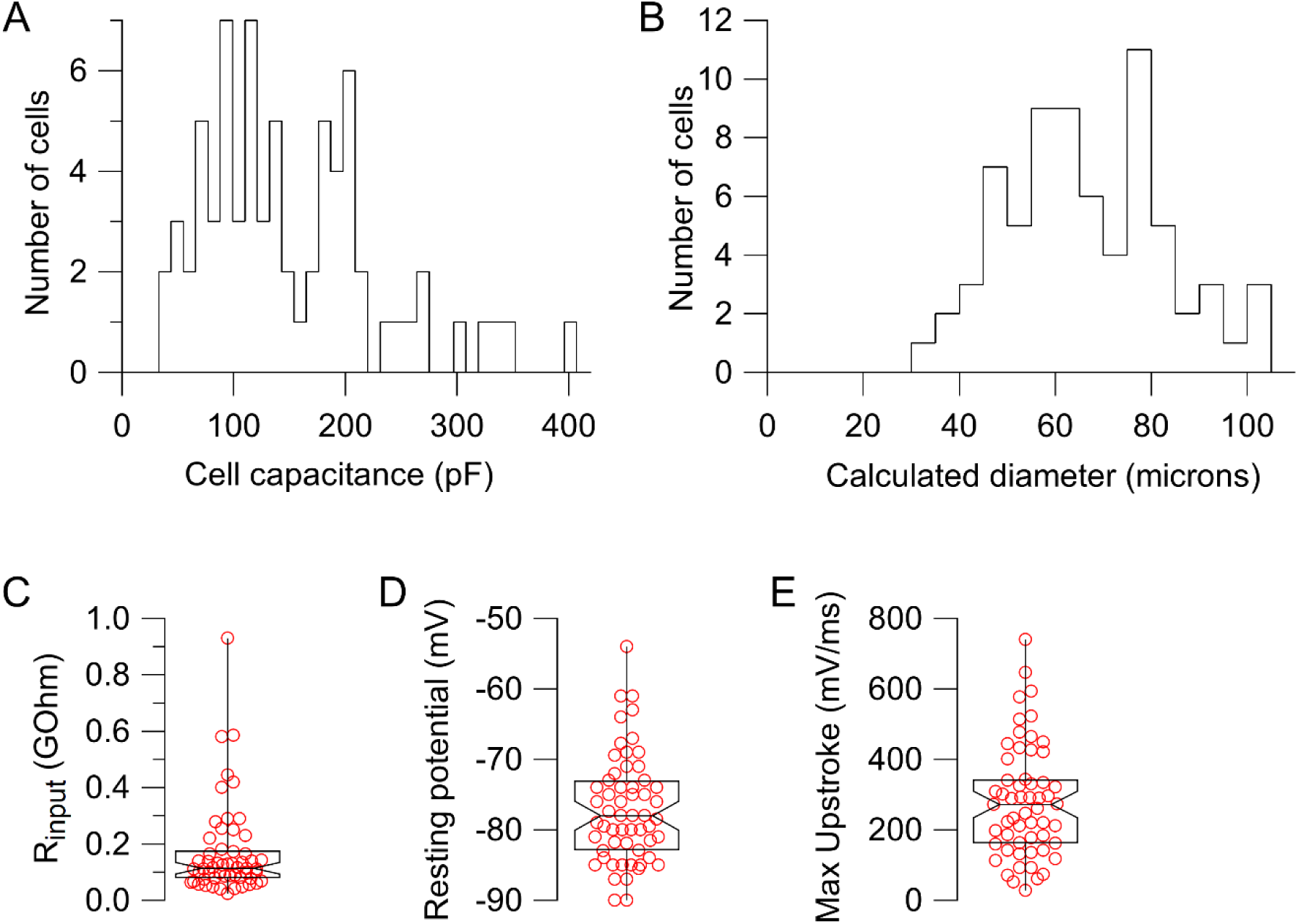
Distribution of cell sizes, input resistance, resting potential, and action potential upstroke velocity of neurons used for experiments (all capsaicin-sensitive). A, Histogram of cell capacitances (n=72). B, Histogram of approximate cell diameters calculated from cell capacitance assuming a spherical cell and specific membrane capacitance of 1 μF/cm^2^ (n=72). C, Tukey box plot of collected data for input resistance, calculated from step from-90 to-85 mV immediately after establishing whole cell recording (22°C, before moving cell to heated solution, n=58). Middle bar shows median, with notch indicating 95% confidence interval for median, lower bar indicates 25% percentile, and upper bar indicates 75% percentile. D, Tukey box plot of collected data for resting potential (37°C, n=57). E, Tukey box plot of collected data for maximal upstroke velocity of the action potential (37°C, n=57).

**Figure S2.**
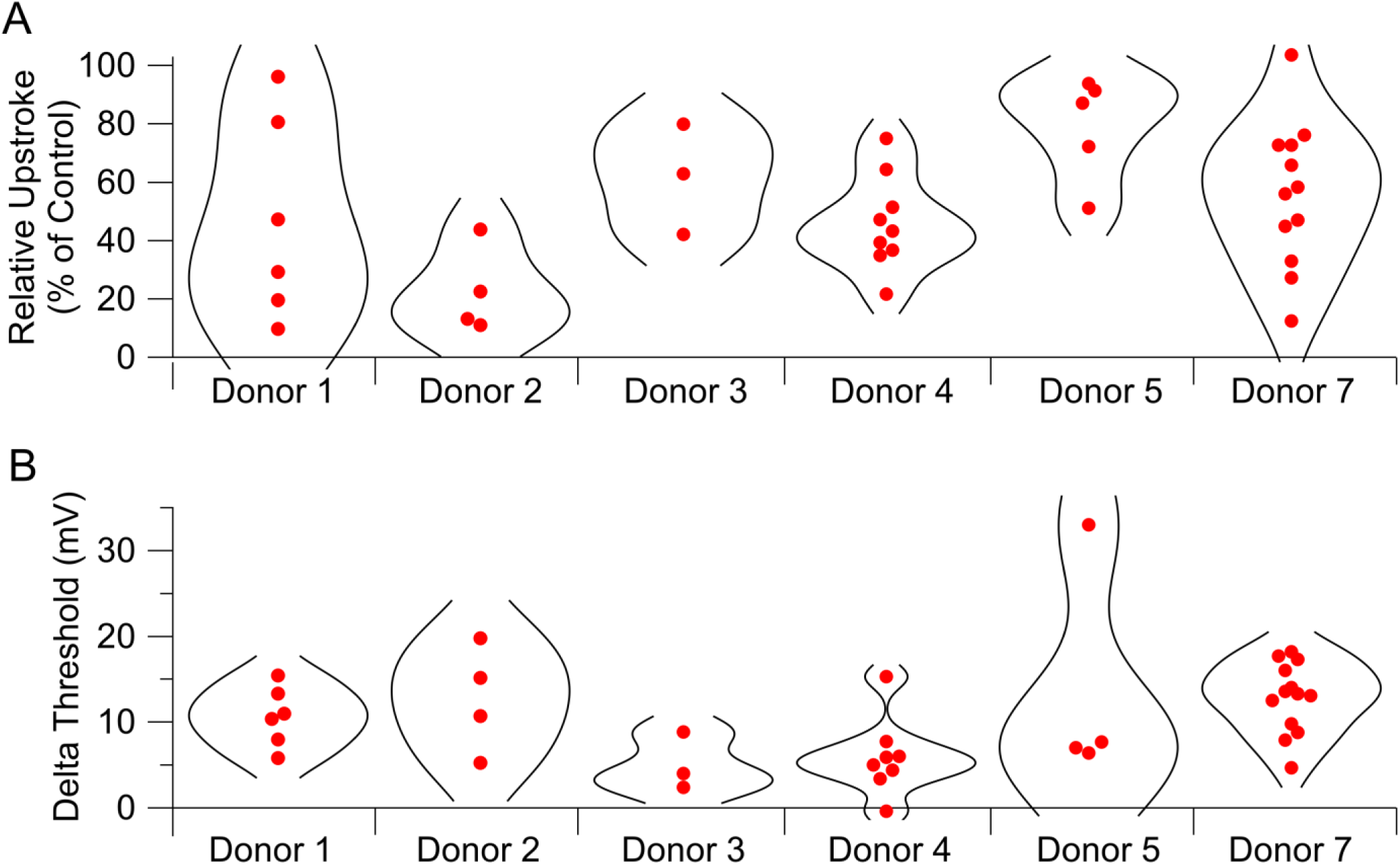
**Effects of 200 nM AM-2099 on action potential parameters compared in cells from individual donors.**

**Figure S3.**
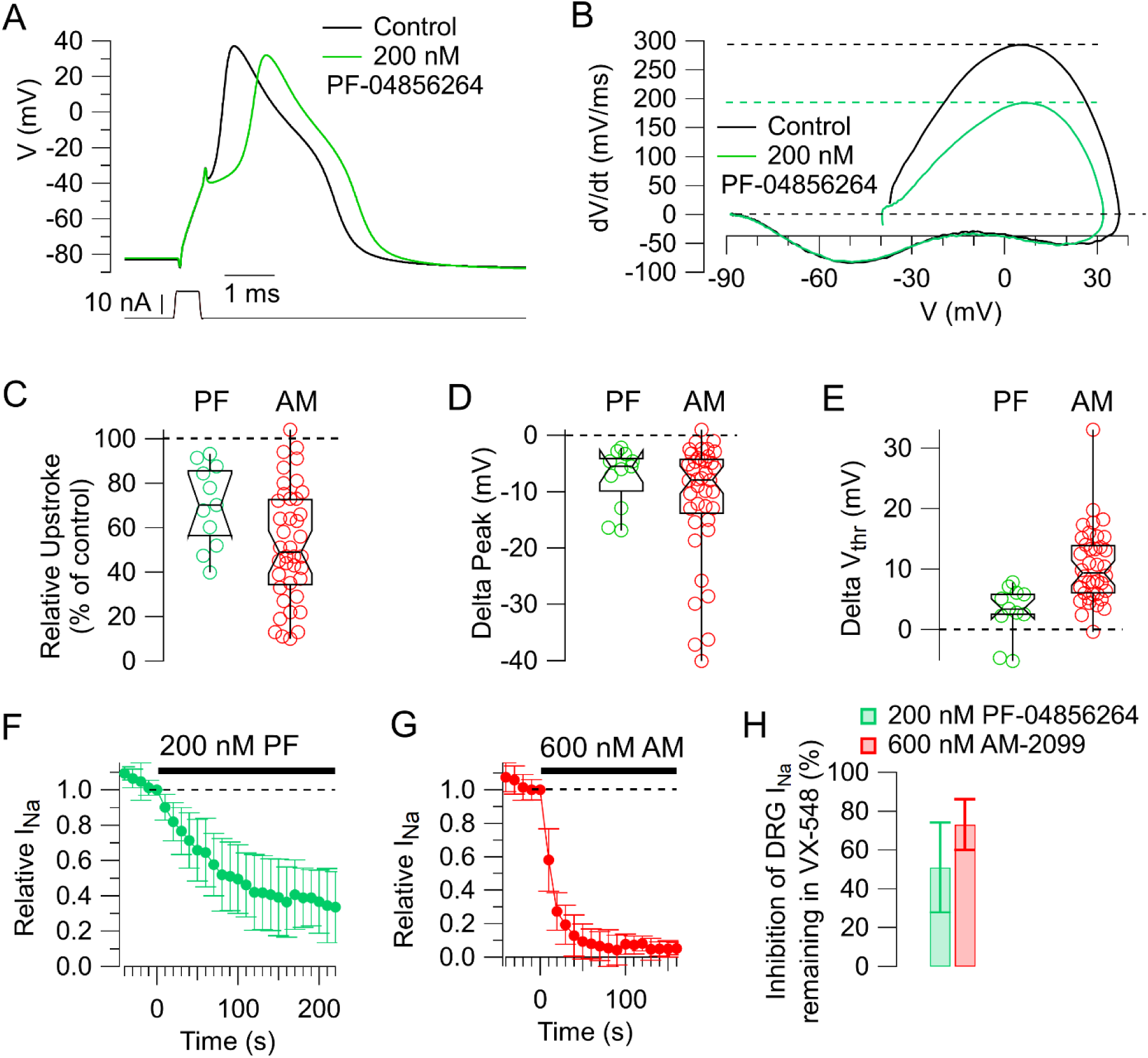
Effects of 200 nM PF-04856264 compared to 600 nM AM-2099. A, Effect of 200 nM PF-04856264 on the action potential in a human capsaicin-sensitive DRG neuron. B, Phase-plane plot of dV/dt versus V for the action potentials in A showing reduction of maximum upstroke velocity and peak. C-E, Collected effects of 200 nM PF-04856264 on action potential parameters (green symbols, n=11) compared to those of 600 nM AM-2099 (red symbols, replotted from Figures 3 and 4). Middle bar shows median, with notch indicating 95% confidence interval for median, lower bar indicates 25% percentile, and upper bar indicates 75% percentile. F, Time-course of inhibition of sodium current from cloned human Nav1.7 channels by 200 nM PF-04856264. Manual patch clamp, 37°C. Mean ± SD. N=7 to 150 s, 6 to 160 s, 5 to 220 s. G, Same for inhibition by 600 nM AM-2099. N=8 to 70 s, 7 to 90 s, 5 to 120 s, 4 to 160 s. H, Effect of 200 nM PF-04856264 on sodium current remaining in the presence of 10 nM VX-548 in human capsaicin-sensitive DRG neurons (green, mean ± SD, n=9) compared to 600 nM AM-2099 (red, re-plotted from Fig 2).

## Notes

### Competing Interest Statement

The authors have declared no competing interest.

### Summary of Updates

Missing number for IC50 of AM-2099, bottom of page 4

